# Statistical validation verifies that enantiomorphic states of cell chirality are determinant dictating the left- or right-handed direction of the hindgut rotation in *Drosophila*

**DOI:** 10.1101/2020.10.08.316182

**Authors:** Tomoki Ishibashi, Mikiko Inaki, Kenji Matsuno

## Abstract

In the left-right (LR) asymmetric development of invertebrates, chirality of cells (cell chirality) plays crucial roles. Left- or right-handed structure of cells consequently directs the morphogenesis with corresponding LR asymmetry. In *Drosophila*, it has been suggested that cell chirality drives the LR-asymmetric development of various organs including the embryonic hindgut. However, this hypothesis is supported only by apparent concordance between these two events and by computer simulations connecting them ^[1–5]^. Therefore, here, we mathematically evaluated the causal relationship between the cell chirality of the hindgut epithelial cells and the LR-directional rotation of the hindgut, which was previously postulated. Our logistic model obtained from various genetic backgrounds significantly explained the correlation between the enantiomorphic states of cell chirality and the LR directionality of hindgut rotation. This model also significantly explained the correlation between cell chirality stochastically formed in advance in each living embryo and the LR-directionality of the following rotation, suggesting the irrelevance of modes how cell chirality is formed. This analysis also showed that the cell chirality is not only sufficient but also required for the rotation. Considering the chronological order of these events, our study validated that cell chirality causally defines the LR asymmetry of the hindgut rotation.

## 1. Introduction

Left-right (LR) asymmetry is often integrated into the basic body plan in bilaterians. For example, internal organs of bilaterians are often LR asymmetric in their morphology and functions. The mechanisms of LR asymmetric development have been well studied in vertebrates ^[6–10]^. For example, in mice, motile cilia locating in the node rotates clockwise and induces a leftward flow of extra-embryonic fluid, which first breaks the LR symmetry ^[11]^. This phenomenon is called as “nodal flow” that subsequently induces the left-side-specific expression of genes, such as *Nodal* and *Lefty*, required for the LR asymmetric development of organs ^[11–13]^. The mechanisms of the LR symmetry breaking are evolutionarily diverged even among vertebrates. For example, in reptile and chick, the nodal flow-independent mechanisms of the LR-symmetry breaking were reported ^[14,15]^. In invertebrates, the chirality of cells (cell chirality) contributes to their LR-asymmetric development ^[16–18]^. An object, such as a cell, is chiral if it cannot be superimposed onto its mirror image. In snails and nematodes, the cell chirality of blastomeres at the initial stage of cleavage determines the subsequent LR-asymmetric arrangement of them, which consequently directs the LR asymmetry of their whole bodies ^[19,20]^.

*Drosophila* has various organs showing stereotypic LR asymmetry, which include, for example, the embryonic gut, male genitalia, and brain ^[21–29]^. The embryonic hindgut is the first organ that shows LR asymmetry during the development of this species ^[21]^. The hindgut is formed in a bilaterally symmetric structure whose anterior part curves toward the ventral side of the embryo and subsequently rotates counterclockwise 90°, which makes it bent rightward ^[21]^. It was shown that the active force for driving the hindgut rotation is generated by the hindgut epithelial tube ^[2,30]^. Before the rotation, apical cell-boundaries of the hindgut epithelial cells slant leftward with respect to the anterior-posterior axis of this organ ^[1–3,31]^. Considering the apical-basal polarity of the hindgut epithelial cells, the three-dimensional structure of them has cell chirality ^[1]^. During the rotation, the cell chirality eventually dissolved so that the cell shape becomes bilaterally symmetric after the rotation of the hindgut ^[1]^. Cooperative studies between computer simulations involving vertex models and image analysis of the hindgut epithelium suggested that the dissolution of the cell chirality induces “cell sliding,” which leads to the LR-asymmetric rotation of the hindgut epithelial tube ^[1,2]^. In cell sliding, the epithelial cells LR-asymmetrically changes their relative position without cell-junctional remodeling, which is coupled with the tube rotation to the corresponding LR directions ^[2]^. Contributions of cell chirality to the LR-asymmetric development were reported in other LR asymmetric organs in *Drosophila*, suggesting that cell chirality has general roles in the LR-asymmetric development of this species ^[4,25,32,33]^. Not only in snails, nematodes, and *Drosophila* but also in Chordata, such as chicken and Larvaceans, the cell chirality was observed and suggested to be involved in the LR asymmetric development of these species ^[34,35]^. In addition to the cell chirality observed *in vivo* during development, various cultured cells from *Dictyostelium* to humans demonstrate intrinsic chirality in their structure and movement ^[36–40]^. Thus, cell chirality is a widely observed property shared among eukaryotic cells.

Using *Drosophila* as a model system, genetic mechanisms underlying the formation of cell chirality has been studied ^[1,31,33]^. *MyosinID* (*MyoID*), a Myosin I family gene and also called *Myosin31DF*, has an activity to induce dextral cell chirality, so that the loss-of-function mutations of *MyoID* resulted in the sinistral cell chirality and the LR-inversion of various organs demonstrating dextral LR asymmetry in wild-type, including the embryonic hindgut ^[1–5, 25, 31, 33]^. *MyosinIC* (*MyoIC*), also called *Myosin61F* and belonging to Myosin I family, bears an activity to induce sinistral cell chirality, which leads to the LR inversion of various organs with handedness upon its overexpression^[5, 22, 30, 41]^. Besides these Myosin I family genes, we previously found that the loss-of-function mutations of *extra macrochaetae* (*emc*) gene, the *Drosophila* ortholog of *Id*, show the randomization of the hindgut handedness ^[31]^. Id family genes encode class V helix-loop-helix (HLH) proteins that inhibit E-box proteins, a class of basic HLH transcriptional factors such as Daughterless (Da) in *Drosophila*, by forming heterodimer, which regulates the balance between cell differentiation and cell proliferation ^[42]^. We found the *MyoID* functions downstream of or parallel to *emc* in the cell chirality formation ^[31]^. However, although genetic mechanisms of cell chirality formation began to be understood, the causal relationship between the cell chirality and hindgut rotation was predicted only on apparent concordance between the enantiomorphic states of cell chirality and the directions of the hindgut rotation in various mutants and on the results of computer simulation recapitulating the rotation of the model gut tube driven by the dissolution of cell chirality ^[1, 2]^. Therefore, the validity of such a cause and effect relationship was still needed.

Here, we mathematically evaluated the causal relationship between the cell chirality and hindgut rotation. We developed a new live-imaging procedure to analyze the cell chirality and subsequent rotation of the hindgut, which provides an opportunity to test the cause and effect relationship between them. Our study validated that the enantiomorphic states of cell chirality causally defines the LR asymmetry of the hindgut rotation.

## 2. Materials and Methods

### 2.1 Fly lines

*Canton-S* was used as wild-type. The following mutants were used: *emc^AP6^*, an amorphic allele (Bloomington #36544) ^[43]^; *da^10^*, an amorphic allele (Bloomington #5531) ^[44]^; and *MyoID^K2^*, an amorphic allele ^[45]^. The following UAS lines were used: *UAS-emc::GFP* ^[46]^; *UAS-da* (Bloomington #51669) ^[47]^; *UAS-MyoIC* ^[22]^; *UAS-MyoID::mRFP* ^[3]^; and *UAS-myr::GFP* ^[48]^. The following *Gal4-driver* lines were used: *NP2432* (Kyoto DGRC #104201) ^[49]^ for the analysis of fixed cell chirality index, and *byn-Gal4* ^[22]^ for the analysis of live cell chirality index. Mutations on the second chromosome and third chromosome were balanced with *CyO, P{en1}wg^en11^* and *TM3, P{GAL4-twi.G}2.3, P{UAS- 2xEGFP}AH2.3, Sb^1^ Ser^1^*, respectively. All genetic crosses were carried out at 25°C on a standard *Drosophila* culture medium.

### 2.2 Live Imaging and laterality score

To visualize the cell membrane of the hindgut epithelium, the expression of *UAS-myr::GFP* was driven by *byn-Gal4* using GAL4/UAS system ^[50]^. *Drosophila* embryos were dechorionated and placed on grape juice agar plates. Embryos of the appropriate genotypes were selected at early stage 12 under a fluorescence microscope and mounted dorsal side up on double sticky tape on slide glasses. These embryos were overlaid by oxygen-permeable Halocarbon oil 27 (Sigma, USA) and a coverslip with spacers of 0.17-0.25 mm-thick coverslips. The live images of apical cell-boundaries in hindgut epithelium were obtained with a scanning laser confocal microscope, LSM 880 (Carl Zeiss, Germany) before the onset of the hindgut rotation. After the imaging, the embryos were continuously cultured at 25°C for 3 hours and the rotational directions of the hindgut were evaluated. As laterality score, the hindgut rotating to the counter-clockwise and clockwise directions was scored 1 and 0, respectively. When the hindgut did not rotate, the laterality score was marked as 0.5.

### 2.3 Analysis of cell chirality index

Cell chirality was analyzed as previously described ^[31]^. Briefly, we obtained the images of apical cell boundaries, which were detected by Myr::GFP signal, in the dorsal part of the embryonic hindgut just before its rotation (at stage 12) using a confocal microscope (LSM880, Carl Zeiss, Germany). Based on these images, the angles (*θ*) between the anterior-posterior axis of the hindgut tube and each cell boundary were determined by ImageJ Fiji v.2.0.0 ^t51]^. To quantify the LR asymmetric slanting of the apical cell boundaries in the hindgut epithelial cells, a cell chirality index was calculated as (*N_R_* — *N_L_*)/(*N_R_*+*N_L_*), where *N_R_* was the number of boundaries with 0° < *θ* < 90° and –180° <–90°, and *N_L_* was the number of boundaries with –90° < *θ* < 0° and 90° < 0 < 180°, for each embryo. The chirality index was measured in a double-blind manner, in which the person analyzing the cell boundaries did not know the genotypes of the embryos and the prospective rotational direction of the hindgut.

### 2.4 Statistical analysis

To test the correlation between the mean cell chirality and the mean of the laterality score in fixed embryos or between the live chirality index and the laterality score in each living embryo, logistic models were used ^[52]^. As a null hypothesis, naive models were used ^[52]^. For the fixed embryos, the mean of laterality score was treated as a response variable, and the mean of the cell chirality index was treated as an explanatory variable. For the live embryos, the laterality score of each individual embryo (0, 0.5, or 1) was treated as a response variable, and the live cell chirality index of each was treated as an explanatory variable. All statistical analyses were performed using R software v.3.6.3 ^[53]^. Graphs were prepared using Matplotlib v.3.0.3 in Python v.3.6.4 ^[54,55]^.

## 3. Results

### 3.1 A logistic model significantly explained the correlation between the enantiomorphic states of cell chirality and the LR directionality of hindgut rotation

The *Drosophila* hindgut is first formed as a bilaterally symmetric invagination whose anterior part curves ventrally at early stage 12 (Figure 1A most left panel) ^[22]^. Then, it gradually rotates counterclockwise 90° as a view from the posterior (Figure 1A, middle two panels). Consequently, this rotation makes the hindgut curved towards the right at the end of stage 13 (Figure 1A, most right panel). The hindgut epithelium has a typical apical-basal polarity, and the inner surface of the hindgut corresponds to the apical surface. Before the rotation of the hindgut, cell chirality of the hindgut epithelial cells is detected in the shape of their apical surface, as the frequency of cell boundaries that slant left or right with respect to the anterior-posterior axis of the hindgut tube shows deviations according to the genetic conditions ^[31]^. For example, cell-boundaries slanting to the left appeared more often than that of the right in wild-type (Figure 1B, left). However, such LR bias found in the angle of cell boundaries is dissolved after the completion of the hindgut rotation (Figure 1B, right). To quantitatively analyze the cell chirality, “chirality index” was formulated previously ^[31]^. The apical cell boundaries were visualized, and the origin of two-dimensional coordinate corresponding the anterior-posterior and left-right axes are placed on a vertex (Figure 1C most left). The cell boundaries exist in the first and third quadrants (shown in orange) and the second and fourth quadrants (shown in turquoise) are defied as Right (orange lines) and Left (turquoise lines), respectively (Figure 1C left and middle). Then, the total numbers of Right and Left were represented by N-Right (*N_R_*) and N-Left (*N_L_*), respectively (Figure 1C, right). The average values of the cell chirality index obtained from embryos (numbers are indicated in Figure 2B as *N_x_*) with each genotype were referred to as “mean cell chirality index.”

**Figure 1.**
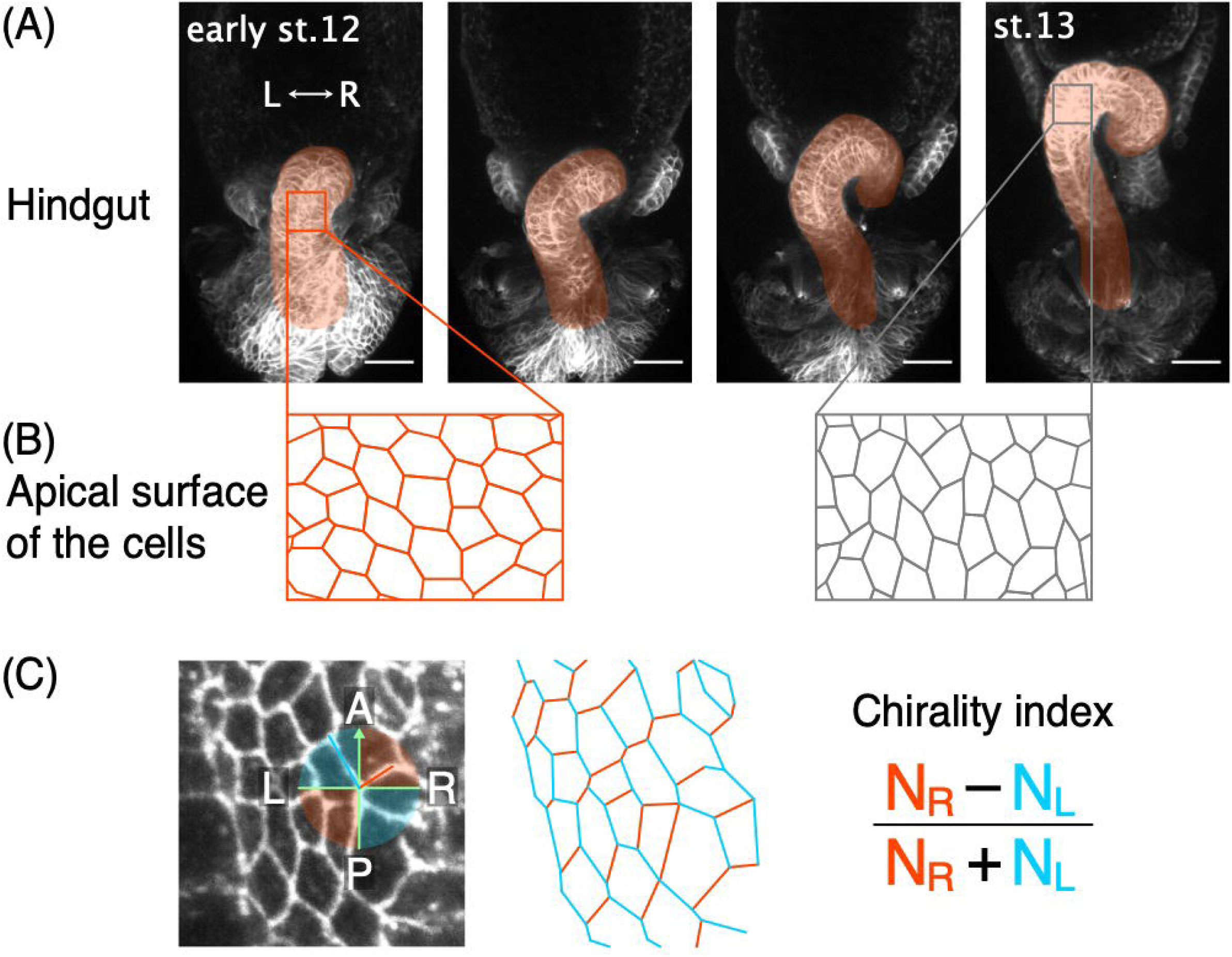
The wild-type *Drosophila* embryonic hindgut rotates 90° counter-clockwise. (A) The *Drosophila* embryonic hindgut (orange) visualized by the expression of *UAS-myr-GFP*, encoding a cell-membrane marker, driven by *byn-Gal4*. The hindgut has a bilaterally symmetric structure whose anterior part curves ventrally at early stage 12. In the wild-type, the hindgut subsequently rotates counter-clockwise and this rotation consequently makes this organ curved to the right. At the end of stage 13, the hindgut exhibits a rightward pointing hook shape. L and R indicate left and right, respectively. (B) Before the hindgut rotation (orange), the apical cell-boundaries of the wild-type hindgut epithelial cells tend to slant to the left, which is defined as the dextral cell chirality. After the hindgut rotation (gray), the cell chirality disappears so that these cells become LR symmetric. (C) Schemas summarizing the procedure used to calculate the cell chirality index. The angle between each boundary and the anterior-posterior (AP) axis of the hindgut was determined. LR represents the left-right axis of the hindgut. Each boundary was classified as a right-tilted (orange) or left-tilted boundary (turquoise). The numbers of right-tilted (*N_R_*) and left-tilted (*N_L_*) boundaries were counted, and the cell chirality index was calculated using the formula shown at the right side of the figure. Scale bars in A are 0.25 *μ*m.

**Figure 2.**
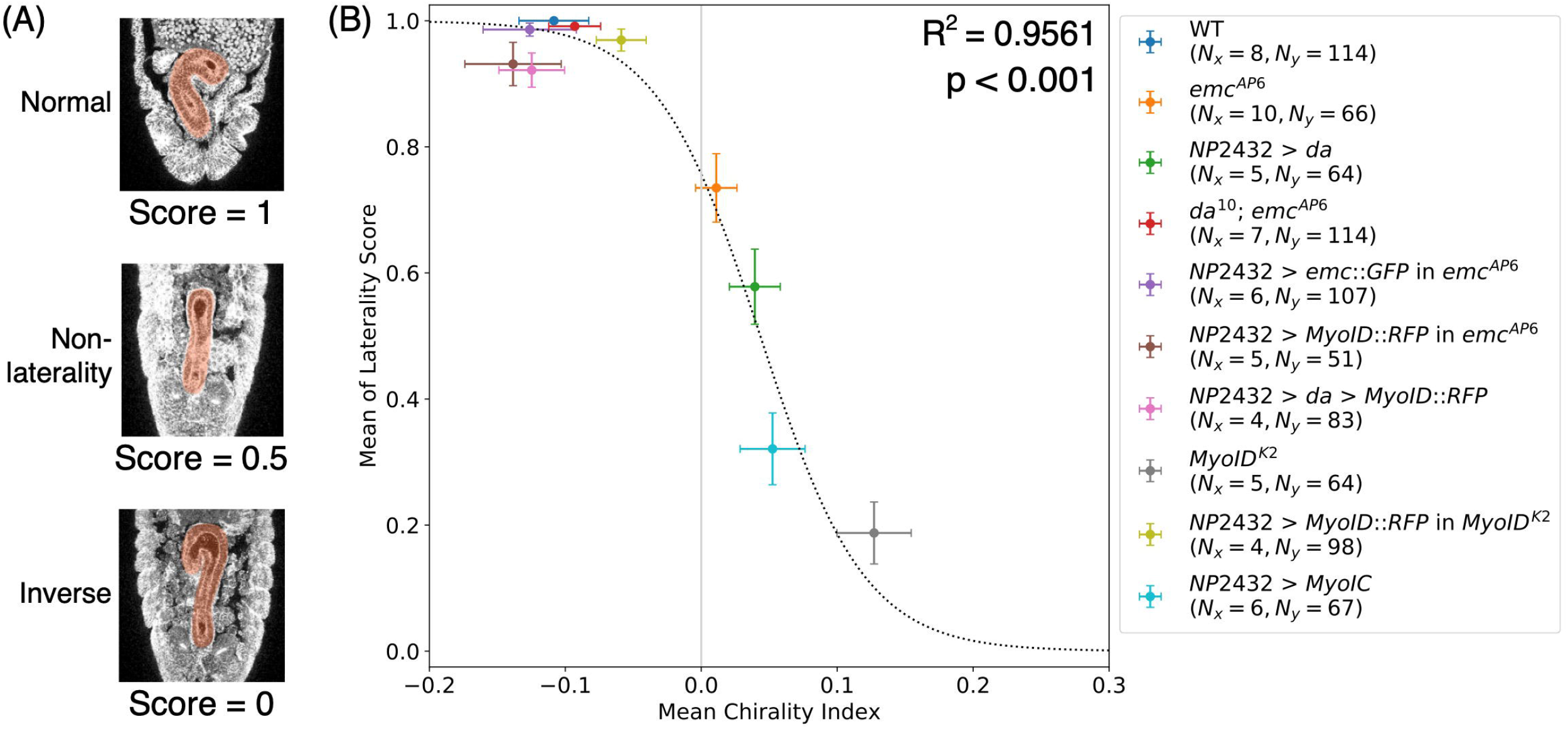
Cell chirality significantly explains the hindgut LR asymmetry of the fixed embryos with various genotype. (A) Typical examples of the hindgut (shown in orange) demonstrating normal LR asymmetry (Normal), bilateral symmetry (non-laterality), and inversed LR asymmetry (Inverse), which was scored as 1, 0.5, and 0, respectively. (B) Graph showing the correlation between the mean cell chirality and the mean value of the laterality score. Some of the data were adopted from our previous paper ^[31]^. The genotypes are shown in the right panel. *N_x_* and *N_y_* indicate the numbers of embryos used for the analyses of the mean chirality index and the mean of laterality score, respectively. The mean values (circles) and standard errors (colored lines) are shown. Dotted line indicates the logistic model that most fits for the data. The coefficient of determination is shown as *R*^2^. The *p*-value of two-tailed t-test was less than 0.001.

We previously showed that the mean cell chirality index strongly correlated with the ratio of the normal hindgut laterality in embryos with various genotypes ^[31]^. To investigate a causal association between the enantiomorphic states of cell chirality and the LR directions of the hindgut rotation, we here analyzed the mathematical characteristics of the correlation between the mean cell chirality index and the rotational direction of the hindgut. We scored the rotational direction of the hindgut in each embryo as a “laterality score”, in which the hindgut of normal LR asymmetry, non-laterality (bilateral symmetry), and inverse LR asymmetry was scored 1, 0.5, and 0, respectively (Figure 2A). The mean of the laterality score was calculated in embryos with each genotype (Figure 2B). Besides the eight genetic conditions analyzed in the previous study, we examined two additions to study the correlation between the mean cell chirality index and the mean of the laterality score (Figure 2B) ^[31]^. As the newly added conditions, double mutant embryos of *da^10^* and *emc^AP6^* (*da^10^; emc^AP6^*) were analyzed (Figure 2B). Our previous analysis revealed that *da* mutation suppresses the LR defects of the hindgut in *emc* mutant because *emc* mutant induces the LR defects through the hyper-activation of *da* ^[31]^. Additionally, embryos overexpressing *UAS-MyoIC*, which has the sinistral activity of the LR asymmetric development, in the hindgut epithelium were examined (Figure 2B) ^[5,30,41]^.

We fitted the logistic model (dotted line) with the chirality index as an explanatory valuable and the mean of the laterality score as an objective variable (Figure 2B). The chirality index significantly explained the mean of laterality score (Quasibinomial generalized linear model (GLM), two-tailed t-test, *p* < 0.001) (Figure 2B, Table S1).

### 3.2 Cell chirality index predicts the LR directions of future hindgut rotation in live embryos

Considering that the cell chirality appears before the initiation of the hindgut rotation, the significant correlation between the mean cell chirality index and the mean of laterality score suggested that cell chirality index of each embryo may determine the LR directionality of the hindgut rotation in an individual. In this case, the chirality index in each live embryo should predict the left or right directionality of the future hindgut rotation afterward. To test this possibility, we developed a new procedure to obtain the cell chirality index in the hindgut of live embryos and then determine the subsequent LR directionality of the hindgut rotation. In this system, the apical cell boundaries of the hindgut epithelium were visualized in live embryos at early stage 12 under a confocal laser microscope, and these embryos were continuously cultured another three hours (Figure 3A). At that point, the rotation of the hindgut was completed under these conditions in wild-type (Figure 3A right). The cell chirality index obtained from this procedure was designated as “live cell chirality index,” and the laterality score of each embryo was marked 1, 0.5, and 0 as the hindgut subsequently showed normal LR asymmetry, non-laterality (bilateral symmetry), and inverse LR asymmetry, respectively.

**Figure 3.**
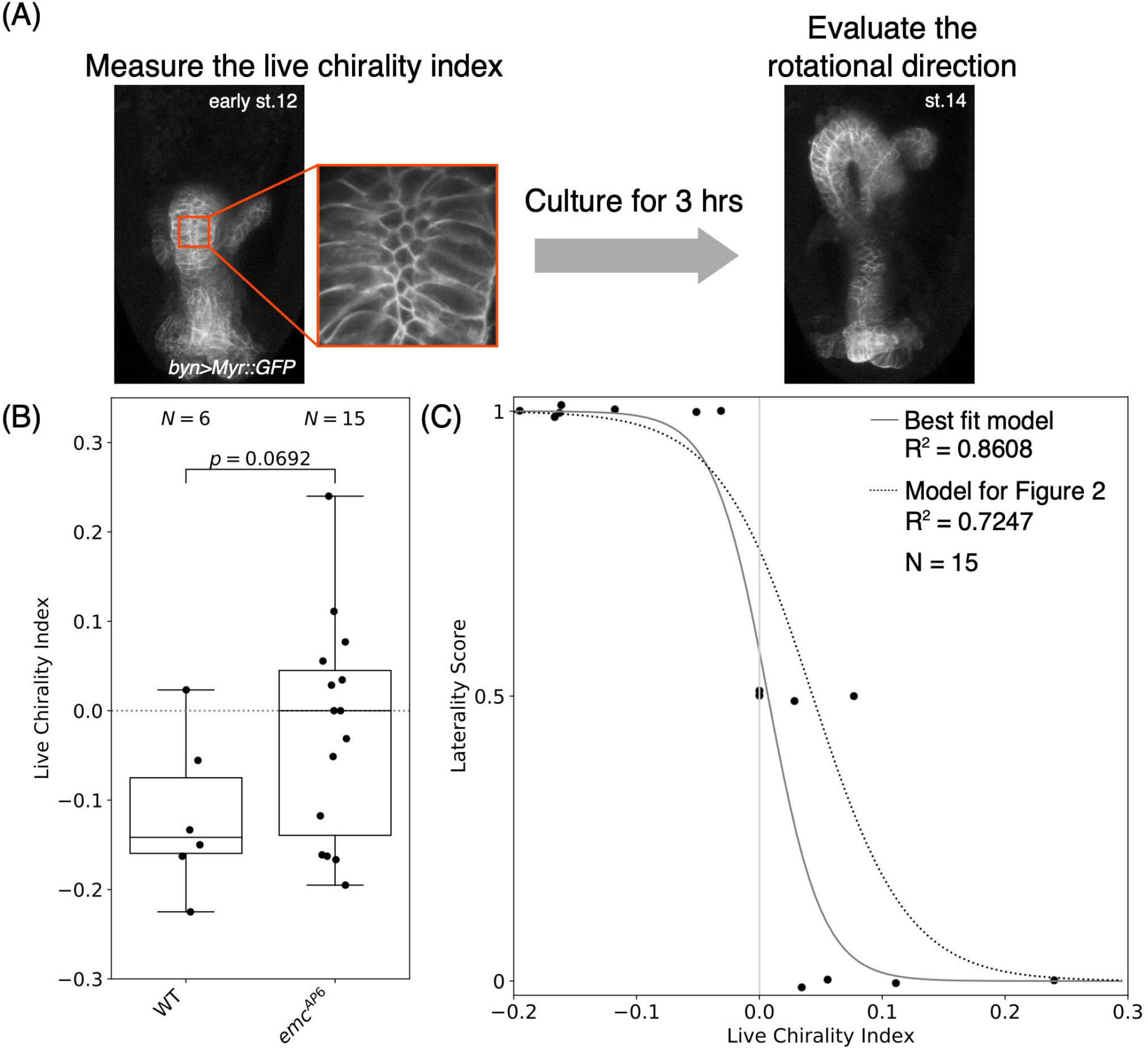
Live cell chirality predicts the rotational direction of the hindgut. (A) Typical images of the hindgut in live embryos. The live images of cell-boundaries visualized by *UAS-myr::GFP* driven by *byn-Gal4* in the hindgut epithelium is shown. At early stage 12, the live cell chirality index of the living hindgut epithelial cells was calculated based on a high magnification image surrounded by the orange square (left panel). After 3 hours of incubation, the laterality score was evaluated as normal LR asymmetry, bilateral symmetry, and inversed LR asymmetry were 1, 0.5, and 0, respectively. (B) Box plot of the live chirality index of the hindgut in the wild-type and *emc^AP6^* mutant embryos. *N* indicates the number of total embryos analyzed. The *p*-value of two-tailed t-test is shown. (C) Graph showing the correlation between the live cell chirality and the laterality score of the hindgut in *emc* mutant. Solid line indicates the best fitted logistic model (Table S2). Dotted line indicates the logistic model obtained from fixed embryos in Figure 2. The coefficient of determination of each model is shown as *R*^2^. *N* indicates the number of total embryos analyzed.

To investigate whether the live cell chirality index correlates with the rotational direction of the hindgut, we found that the LR-randomization phenotypes of *emc^AP6^* homozygote were appropriate for this purpose. Only in *emc^AP6^* mutant among the ten different mutants examined here, the mean cell chirality index was nearly zero with deviation ranging from plus to minus values (Figure 2B). In addition, the hindgut of these embryos shows normal and inverse LR asymmetry as well as non-laterality ^[31]^. Thus, we speculated that each individual embryo has stochastically fluctuated cell chirality, which may induce the hindgut rotation to the direction corresponding to the cell chirality index of each. Based on this assumption, we evaluated the consistency between the chirality indexes obtained from fixed and live embryos, we analyzed the mean of live chirality indexes in wild-type and *emc^AP6^* homozygous embryos. The mean live chirality indexes of wild-type and *emc^AP6^* homozygous embryos were −0.12 ± 0.04 and −0.02 ± 0.03, respectively, which were very similar to those of fixed embryos as reported before (wild-type, −0.11 ± 0.03; *emc^AP6^* homozygote, 0.01 ± 0.02), although these mean values of live cell chirality indexes were not significantly different in this experiment (Two-tailed t-test, *p* = 0.069217) ^[31]^. We also found that each individual embryo of *emc^AP6^* homozygote showed a broad range of live chirality indexes, which were distributed from plus to minus values, as compared with that of wild-type (Figure 3B).

Given that the values of the live chirality index agreed well with those of the cell chirality index in the fixed embryos, we next conducted a statistical correlation analysis between the live cell chirality index and the laterality score of each individual *emc^AP6^* mutant embryo. We fitted the logistic model (solid line) with the live chirality index as an explanatory variable and the laterality score of each embryo as an objective variable and found that live cell chirality, which was determined prior to the onset of the hindgut rotational direction significantly explained the rotational directions of the hindgut (Quasibinomial GLM, two-tailed t-test, *p* < 0.01; Figure 3C; Table S2). Importantly, the logistic model (dotted line) obtained the fixed embryos with various genetic conditions as described in Figure 2 significantly fitted the data obtained from each living *emc^AP6^* mutant embryo (Quasibinomial GLM, likelihood ratio test, 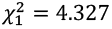, *p* < 0.05), and it explained 72.5% of the total variation in the rotational direction of live *emc^AP6^* mutant hindgut (Table S3). These results suggested that the cell chirality index correlates with the rotational direction of the hindgut, disregarding that the enantiomorphic states of cell chirality are determined by genetic conditions or stochastic fluctuations. Although, with just a statistical correlation, we could not conclude whether cell chirality is the cause or effect. Nevertheless, considering the clear chronological sequence of the cell chirality formation and the initiation of the hindgut rotation, we concluded that the enantiomorphic states of cell chirality direct the LR asymmetry of the hindgut rotation.

## 4. Discussion

In previous studies, it was suggested that the enantiomorphic states of cell chirality in the hindgut epithelium determine the directionality of the hindgut rotation ^[1, 2, 31]^. Mainly, dextral and sinistral cell chirality on average lead to the counter-clockwise and clockwise rotation of the hindgut, respectively ^[1, 2]^. This postulation was firstly supported by the observations that the mean states of cell chirality and the LR directions of the hindgut rotation correlate to each other in various mutants ^[1, 2, 31]^. For example, in *MyoID* mutants where cell chirality becomes sinistral, their hindgut rotates clockwise, whereas wild-type embryos demonstrate dextral cell chirality and the counter-clockwise rotation of the hindgut^[1]^. Additionally, the cell chirality vanishes on average in *Drosophila E-cadherin* or *emc* mutants in which the LR-directions of hindgut rotation are randomized or the hindgut remains LR symmetric ^[1, 31]^. Secondary, computer simulation employing vertex models suggested that the enantiomorphic states of cell chirality initially introduced into the model defines the LR direction of the model-gut tube rotation and successively drives the tube rotation to the direction determined at the outset ^[1]^. Although these evidences consistently support the postulation, the hypothesis has been grounded on the apparent concordance between the states of cell chirality and the LR directions of the hindgut rotation and on the prediction from computer simulations ^[1–2]^. Thus, the cause and effect relationship between them needed to be verified.

To address this issue, here, we mathematically evaluated this hypothetical causal relationship by analyzing the correlation between the enantiomorphic states of cell chirality and the LR direction of the hindgut rotation. The means of the cell chirality index and laterality score in the fixed embryos were obtained for each of ten different genetic conditions, and the correlation between them was analyzed. In our logistic model, the mean cell chirality index representing each genotype as an explanatory variable significantly fitted to the mean of the laterality score (Figure 2B). Therefore, the mechanisms by which the cell chirality defines the LR-directionality of the hindgut rotation may be similar regardless of genetic backgrounds. This result also demonstrates that the LR directionality of the hindgut rotation is statistically explained even only by the enantiomorphic states of cell chirality as the average of embryos with each specific genotype.

In the correlation analysis of the fixed embryos, the mean of cell chirality index was 0.01 ± 0.02 in *emc^AP6^* mutant embryos, nevertheless, they showed the hindgut rotating to the directions of either clockwise or counter-clockwise or staying as bilaterally symmetric shape (Figure 2B) ^[31]^. Thus, one can argue that cell chirality may not be essential for directing the hindgut rotation. Our analyses here addressed this question. In this study, we developed a method to determine the live cell chirality and the directions of the hindgut rotation in each living embryo. We found that the live cell chirality indexes representing each individual embryo of *emc^AP6^* mutant are varied widely among embryos and distributed from plus to minus values, although the mean value of the live cell chirality index of these embryos was nearly zero (Figure 3B). Therefore, this mutant background was found useful to analyze the correlation between the states of cell chirality and the rotational directions of the hindgut in each live embryo. Our logistic model formulated from the data of the live *emc^AP6^* mutant embryos showed a significant correlation between the live cell chirality index and the laterality score of each individual embryo (Figure 3C). Therefore, in each of these embryos, the live cell chirality index significantly explained the rotational directions of the hindgut or its bilateral structure. Based on these results, we ascertained that the cell chirality is not only sufficient but also required for the hindgut rotation because our model predicts that living embryos with zero cell chirality index tend to have bilaterally symmetric hindgut.

Importantly, the logistic model obtained from the fixed embryos with various genetic conditions significantly fitted the data obtained from each living *emc^AP6^* mutant embryo (Quasibinomial GLM, likelihood ratio test, 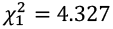, *p* < 0.05), and it explained 72.5% of the total variation in the rotational direction of *emc^AP6^* mutant hindgut (Figure 3C; Table S3). Given that individual embryos all carrying the same *emc^AP6^* mutant background showed a wide range of live cell chirality index, cell chirality may be formed stochastically in each individual under this condition. Nevertheless, the logistic model formulated from the data of fixed embryos with various genotypes significantly explains the data that presumably reflects the stochastic events in each *emc^AP6^* individual. Therefore, underlying mechanisms in which cell chirality drives the LR-directional rotation of the hindgut are common between the cell chirality formed by the modes determined genetically and stochastically.

In this study, we mathematically confirmed the causal relationship between cell chirality and the hindgut rotation. Our logistic model also implied that the cell chirality is sufficient and necessary for the hindgut rotation. Considering the time course of the events in which cell chirality formation precedes the initiation of the LR-directional rotation of the hindgut *in vivo*, our presented results demonstrate that the cell chirality is the primary cause of the hindgut rotation. Since the cell chirality is involved in the LR-asymmetric development of various organs in *Drosophila*, such cause and effect relationship can be applicable to explain their morphogenesis. Furthermore, our idea could be extended to the LR-asymmetric developments in other various species, which are known to involve the cell chirality.

## Supplementary Materials

### Appendix

**Table S1.** Factors affecting the variation in the laterality score of the embryonic hindgut with various genotypes.

|  | *t*<sub>8</sub>-value | *p*-value |
| --- | --- | --- |
| Intercept | -4.042 | 0.003723 |
| Chirality Index | -5.992 | 0.000326 |

**Table S2.** Factors affecting the variation in the rotational direction of the hindgut in *emc^AP6^*.

|  | *t*<sub>13</sub>-value | *p*-value |
| --- | --- | --- |
| Intercept | -1.149 | 0.27109 |
| Live Chirality Index | -3.256 | 0.00625 |

**Table S3.** Logistic regression analyses between the laterality score and live chirality index in *emc^AP6^*.

|  | Model | *R*<sup>2</sup> |
| --- | --- | --- |
| Best fit model | $1 - \frac{1}{1 + e^{-4.531 \times (x - 0.00697)}}$ | 0.8607872 |
| Model fitted for Figure 2 | $1 - \frac{1}{1 + e^{-0.410 \times (x - 3.42)}}$ | 0.7246853 |

## Author Contributions

Conceptualization, T.I. and K.M.; Methodology, T.I.; Investigation, T.I. and M.I.; Data Analysis, T.I.; Writing - Original Draft Preparation, T.I.; Writing - Review & Editing, T.I., M.I., and K.M.; Supervision, K.M.; Funding Acquisition, T.I., M.I., and K.M. All authors have read and agreed to the published version of the manuscript.

## Acknowledgments

We thank N.S. Moon and S. Noselli for fly stocks. We also thank the Bloomington Drosophila Stock Center (Indiana University), the Drosophila Genetic Resource Center (Indiana University), and the Kyoto Stock Center (Kyoto Institute of Technology) for fly stocks, and the Developmental Studies Hybridoma Bank (University of Iowa) for antibodies.

